# Successes and failures of the live-attenuated influenza vaccine, can we do better?

**DOI:** 10.1101/424275

**Authors:** Laura Matrajt, M. Elizabeth Halloran, Rustom Antia

## Abstract

Live-attenuated vaccines are usually highly effective against many acute viral infections. However, the effectiveness of the live attenuated influenza vaccine (LAIV) can vary widely, ranging from 0% effectiveness in some studies done in the United States to 50% in studies done in Europe. The reasons for these discrepancies remain largely unclear. In this paper we use mathematical models to explore how the efficacy of LAIV is affected by the degree of mismatch with the currently circulating influenza strain and interference with pre-existing immunity. The model incorporates two key antigenic distances – the distance between pre-existing immunity and the currently circulating strain as well as the LAIV strain. Our models show that a LAIV that is matched with the currently circulating strain is likely to have only modest efficacy. Our results suggest that the efficacy of the vaccine would be increased (optimized) if, rather than being matched to the circulating strain, it is antigenically slightly further from pre-existing immunity compared with the circulating strain. The models also suggest two regimes in which LAIV that is matched to circulating strains may provide effective protection. The first is in children before they have built immunity from circulating strains. The second is in response to novel strains (such as antigenic shifts) which are at substantial antigenic distance from previously circulating strains. Our models provide an explanation for the variation in vaccine effectiveness, both between children and adults as well as between studies of vaccine effectiveness observed during the 2014-15 influenza season in different countries.

**Significance Statement:** The live-attenuated influenza vaccine, in principle provides an important intervention for the control of both seasonal and pandemic influenza. However vaccine effectiveness studies have found seemingly contradictory results with effectiveness ranging from 0 to 50%. Based on mathematical models we suggest that a major factor responsible for the variable efficacy of the vaccine is negative interference – where pre-existing immunity precludes the vaccine from working. Our models suggest that there are broad regimes for which LAIV will fail to be immunogenic, but also allow us to make suggestions for the choice of vaccine strain that will allow optimization of protective immunity in different scenarios.

---

Seasonal influenza viruses continue to pose significant disease and economic burden, with the Centers for Disease Control and Prevention (CDC) estimating between 12,000 and 56,000 deaths per year, and between 9.2 and 35 million illnesses per year [1, 2] in the United States (US). Due to the constant mutation of influenza viruses, influenza vaccines need to be updated annually, resulting in vaccines with varying degrees of efficacy depending on the currently dominant circulating type (influenza A versus B), subtype (H3N2 vs H1N1) and strain. Currently, there are two types of vaccines against seasonal influenza: inactivated injectable influenza vaccines (IIV) and a live-attenuated influenza vaccine (LAIV), administered as a nasal spray.

Universal influenza vaccination has been recommended in the United States since 2010, with both IIV and LAIV. Because early studies showed that LAIV’s highest efficacy is among young children [3], its use has been concentrated mostly in the younger age groups. Furthermore, several studies done before 2014 comparing the effectiveness of LAIV and IIV found the former to have a much better effectiveness in children than the latter [4–7]. Specificially, Ambrose *et al*. [6] found that children vaccinated with LAIV experienced up to 48% fewer cases than children vaccinated with IIV. Based on these studies, the CDC’s Advisory Committee on Immunization Practices (ACIP) preferentially recommended the use of LAIV in healthy 2 to 8 years old children for the 2014 influenza season. The United Kingdom (UK) started a universal pediatric vaccination program with LAIV in 2013 and, a year later, LAIV was offered in Finland to two year-olds for the first time. However, vaccine effectiveness studies done in the US showed mixed LAIV effectiveness against the corresponding predominant strains during the 2013-2016 influenza seasons [8] ranging from no effectiveness to 17% effectiveness in studies done by the CDC and the Department of Defense [9–13] and 50% effectiveness in a study funded by the manufacturer of LAIV [14]. As a result, the ACIP suspended the recommendation of LAIV for the next two seasons (2016/17 and 2017/18) [15, 16] (the ACIP has since then resumed its recommendation for the 2018/19 season). In contrast, studies done in Canada, the UK and Finland told a very different story, with effectiveness ranging from 31% to 57.6% [17–21].

The reasons for the discrepancies observed in the LAIV effectiveness in different studies and parts of the world remain largely unknown. Some studies suggest that the low effectiveness of the LAIV could be due to a reduction in fitness of the H1N1 component of the vaccine [22], however, the predominant strain in 2014/15 was H3N2. Low and variable efficacy of both LAIV and IIV can be due to mismatches between the vaccine strain and the circulating (challenge) strain (vaccine eliciting an ineffective immune response). Alternatively, low efficacy could be due to a vaccine’s reduced ability to generate an immune response. There are many reasons why a vaccine might not be sufficiently immunogenic [23, 24]: problems might have arisen with the strain contained in the vaccine during production [25], the age of the host (influenza vaccines result in lower immune responses in people over 65 years old [26, 27]) or pre-existing immunity affecting the ability of the vaccine’s immunogenicity [28]. The latter is called negative interference. In a seminal paper in 1999, Smith *et al*. [29] used a mathematical model to provide a quantification of negative interference for repeated influenza vaccination with IIV. They introduced the *antigenic distance* hypothesis, which establishes that “variation in repeat vaccine effectiveness is due to differences in antigenic distances among vaccine strains and between the vaccine strains and the epidemic strain in each outbreak”.

Negative interference has been well documented and studied for IIV, with several studies showing that depending on the year and circulating serotype, people who have been vaccinated in the previous year(s) have a decreased vaccine efficacy compared with those who have not been vaccinated [29–41]. This phenomenon has been less studied for LAIV [34, 38]. Here, we use mathematical models to explore how the antigenic distance hypothesis can be used to explain differences in the vaccine effectiveness of LAIV. In particular, we wish to evaluate how pre-existing immunity, acquired naturally or through vaccination, affects the effectiveness of LAIV. We hypothesize that differences seen in vaccine effectiveness across influenza seasons and across different countries arise from the interplay of three antigenic distances: the distance between the vaccine strain and pre-existing immunity, the distance between the challenge strain and the vaccine strain, and the distance between the challenge strain and pre-existing immunity.

We developed within-host mathematical models describing the interaction between influenza viral infections, vaccination, and the immune response. Our models recapitulate the main features of influenza infection and vaccination. Our models predict broad regimes for which the LAIV would be ineffective, but they also provide guidance as on how to choose the vaccine strain for it to successfully protect the host. Further, our results suggest that under certain conditions, choosing a vaccine antigen different from the challenge strain, and more distant from pre-existing immunity might result in better vaccine protection.

## Results

### Model

Building on [42] we use a simple target-cell limited model for the dynamics of influenza infection outlined in Figure S1. In the SI we show that our results are robust to modifications of the model, including the incorporation of innate immunity. The adaptive immune response is generated by B-cell clones that produce antibodies, with different clones having different affinities to different viruses. The effectiveness of antibody-binding and the stimulation of B-cells depend on a one-dimensional antigenic distance between the virus and the B-cells, measured as a percentage of the difference between the antibody receptors and viruses (full description in Methods). Clones having a higher affinity are stimulated more, and generate larger numbers of B-cells and antibodies. Furthermore these antibodies are more effective at controlling the virus. In our simulations, antibodies stop being cross-reactive once they are 30% different from the virus. Hence, antigenic drift occurs in our models for parameter regimes where the antigenic distance between the challenge virus and previously circulating virus is in that range (20-30%). We modeled preexisting immunity as closest to the virus strain predominantly circulating in the population in the previous year. We define a within-host measure of vaccine effect *υe_ω_*, that quantifies the extent to which the vaccine helps to reduce the viral load of subsequent challenge infections (virus density integrated over the course of the infection as described in the Methods). With a low *υe_ω_*, a vaccinated individual will see little additional benefit compared to that same individual had s/he not been vaccinated when presented with subsequent challenge infections with that particular strain.

### Model recapitulates key features of influenza infection and vaccination

Our model recapitulates key features of primary influenza infection. Figure 1 shows the model dynamics during an influenza primary infection (*i.e*, an infection with a virus for which there is no pre-existing immunity). Viral load peaks around day 3, reaching well-established viral load values [43] (Fig. 1). The B-cell clones closest to the virus are stimulated the most, and the stimulation decays with distance (Fig. 1, pie chart). These clones, in turn, produce elevated levels of antibodies within 2-3 days of stimulation (Fig. 1).

**Figure 1:**
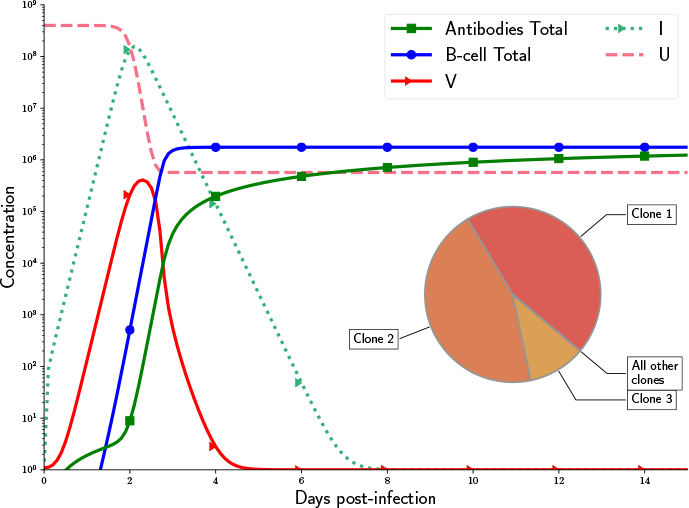
Dynamics of a primary influenza infection for Model 2 (full details in SI). Shown here: uninfected cells *U*, infected cells *I*, virus *V*, total B-cell and antibody dynamics. Pie chart: Contribution of each clone to the total B-cell and Antibody response. The virus *V* was closest to clone 1 in this simulation with the parameters given in Table S1. The model was run with 20 clones.

We now consider how pre-existing immunity affects boosting by a LAIV. If a vaccine strain is closely matched to pre-existing immunity, the model predicts that the pre-existing immunity will control the vaccine and no B-cell clone will be significantly boosted (Fig. 2A) resulting in a poorly immunogenic vaccine. In this case, no new B-cell clone is stimulated but the antibodies matching the primary infection get boosted. However, once a vaccine strain is sufficiently different from pre-existing immunity (~30% for our model parameters), the attenuated virus contained in the vaccine is no longer controlled by pre-existing immunity and this results in a high viral load and in the stimulation of those B-cell clones and antibodies closest to the vaccine strain (Fig. 2B). Interestingly, our model predicts that the antibodies stimulated during the primary infection will always get stimulated (even if this stimulation is very small) during subsequent infections, so that their concentration will always be higher than the other clones, consistent with the Original Antigenic Sin theory [44].

**Figure 2:**
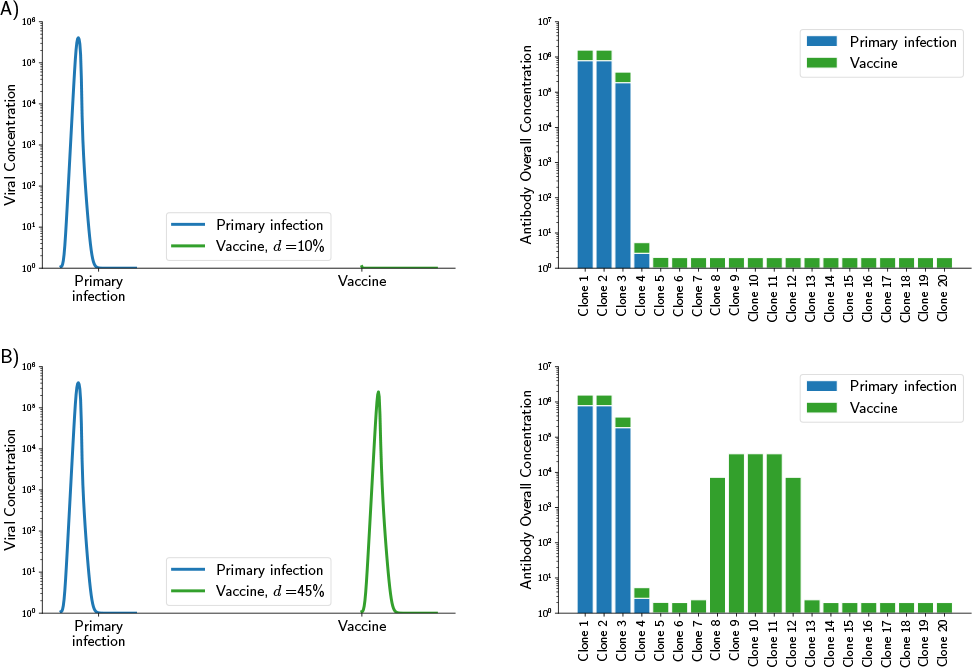
Simulated primary infection and vaccine dynamics for three vaccine strains: A) a slightly different strain, differing only 10% from pre-existing immunity (top row), B) a substantially different strain with no cross-reactivity, 45% difference to pre-existing immunity. For each row, the left panel shows the viral dynamics of primary infection followed by vaccination and the right panel shows the overall antibody concentration. (A) Pre-existing immunity controls the vaccine antigen with no stimulation of any antibody clone, (B) vaccine escapes pre-existing immunity resulting in the stimulation of the B-cell and antibodies closest to the vaccine strain. However, for all cases, antibody clones closest to pre-existing immunity are always boosted in addition to those clones closest to the vaccine.

### Optimal vaccine strain selection

We now address the question of how a vaccine affects the dynamics following challenge with a new strain of influenza. To model prior exposure to influenza we begin by setting the pre-existing level of immunity to that generated by a primary infection as described above. We then consider how the severity of infection with a new virus strain (termed the challenge strain) can be reduced by vaccination. In particular we consider how the severity of infection with the challenge strain depends both on its antigenic distance from pre-existing immunity as well as the antigenic distance of the vaccine from both pre-existing immunity and the challenge strain. We denote the distances between vaccine and pre-existing immunity, vaccine and challenge, and challenge and pre-existing immunity by *d*(*V, P I*), *d*(*V, C*) and *d*(*C, P I*) respectively. In Figure 3A we plot how the severity of an infection with the challenge strain (defined as the AUC of the infection) depends on the distance from the challenge to pre-existing immunity and the distance from vaccine to pre-existing immunity. In Figure 3B we plot *υe_ω_*, the extent to which the vaccine reduces the severity of infection, and in particular how *υe_ω_* depends on the antigenic distance from the vaccine strain to pre-existing immunity and the distance from the challenge to pre-existing immunity.

**Figure 3:**
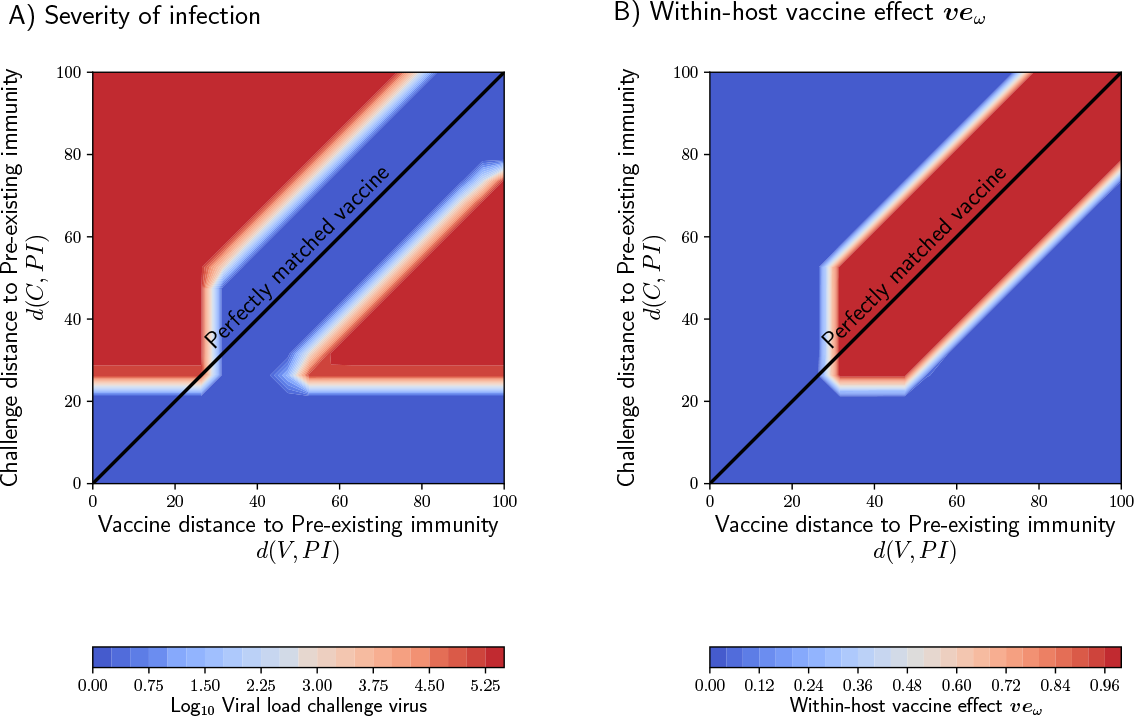
A) Contour plot representing the AUC for the viral load following a challenge infection. Here we used Model 2 (SI) with 20 clones (parameters described in Table S1), where we assumed that an attenuated influenza virus will make the antibody killing rate more efficacious (*k_V_ ac* = 25*k*, see SI for details). B) Contour plot representing the within-host vaccine effect *υe_ω_* as a function of the distance from the vaccine strain to pre-existing immunity and the distance from the challenge to pre-existing immunity. The within-host vaccine effect is a measure of the ability of the vaccine to prevent further infections.

We consider the following parameter regimes for the distance between pre-existing immunity and the challenge strain.

i. When the antigenic distance between pre-existing immunity and the challenge strain is small (*d*(*C, P I*) ≤ 20%). Our model predicts that pre-existing immunity will prevent proliferation of a challenge virus. This can be seen in Fig. 3A where we note that provided *d*(*C, P I*) ≤ 20% the severity of the infection (its AUC) is very low, irrespective of the vaccine. Consequently the vaccine will have a very low *υe_ω_* as seen in Fig. 3B since pre-existing immunity alone is sufficient to control the virus. At the population level, this corresponds to a scenario where the challenge strain is not significantly different from the previous year (a mild year). In this scenario pre-existing immunity prevents infections in most individuals and it also prevents a matched vaccine from boosting the immune response. We also note that unless the vaccine is antigenically much more distant from pre-existing immunity than the challenge strain, pre-existing immunity will control the vaccine (Figs. S2A and B and S3A and B). Regardless the vaccine will have little efficacy against that year’s virus.
ii. We then consider the regime where the antigenic distance between pre-existing immunity and the challenge strain is just a bit too large for pre-existing immunity to immediately clear the challenge strain (20% ≤ *d*(*C, P I*) ≤ 40%). This scenario corresponds to seasonal antigenic drift with the challenge strain causing an infection. The ability of the vaccine to prevent further infections depends on the antigenic distance between the vaccine and pre-existing immunity. If the vaccine is antigenically closer to pre-existing immunity than the challenge strain (region where *d*(*V, P I*) ≤ 20% and to the left of the diagonal black line) then we find that the replication of the vaccine is controlled by pre-existing immunity and it does not boost immunity or affect subsequent infection with the challenge virus (Figs. S2C and S3C). In this case, we expect to see low vaccine effectiveness. The best outcome is when the vaccine is antigenically a little more distant from pre-existing immunity than the challenge strain. In this regime the vaccine boosts immunity that can help control the challenge strain (Figs. S2D and S3D). Analogously, in Fig. 3B we see that the vaccine has optimal within-host vaccine effect when it is slightly more distant from pre-existing immunity than the challenge strain. At the population level, we expect to see a high vaccine effectiveness. When the vaccine is antigenically very distant from both pre-existing immunity and the challenge strain, then the response it elicits does not affect the challenge strain.
iii. Finally we consider a scenario where the challenge strain is antigenically very distant from pre-existing immunity (e.g. *d*(*C, P I*) *>* 40% in our model). This corresponds to a year where there is a major antigenic shift, such as the introduction of the 2009 H1N1 influenza A pandemic strain. In this situation, we see in Fig. 3A and B that the vaccine is most effective when it is antigenically matched to the challenge strain. In this parameter regime, pre-existing immunity has little effect on either vaccine or challenge virus.

### Application to the 2014-2015 influenza season

The 2014-2015 influenza season was characterized as a moderately severe season, with the H3N2 influenza A strain being the dominant circulating strain in North America and Europe [45, 46]. While vaccine effectiveness studies done in the US found contradictory results with one study showing that the LAIV was not efficacious (VE of −3% (CI: −50% to 29%))[13] and the other one showing a vaccine effectiveness of 30% (95% CI, −6% to 54%)[12], studies done in the UK found LAIV to be 35% efficacious (95% CI: −29.9 to 67.5) [18]. Skowronski *et al* [20] computed the antigenic distances between the vaccine, the previous year’s vaccine and the epidemic challenge for the H3N2 strain for the 2014-2015 influenza season. The vaccine was identical to the previous year’s vaccine and they found that the antigenic distance between the vaccine and the epidemic challenge to be 4 (equivalent to 20% in our model), so that *d*(*V, P I*) = 0 and *d*(*V, C*) = 20 = *d*(*P I, C*).

Children in the US have been routinely vaccinated against seasonal influenza since the mid-2000s [47]. The CDC estimates over 58% vaccination coverage of children in the US for the influenza seasons 2013-2017 [48], so it is expected that, during the 2014-15 influenza season, most of the children in the US had some pre-existing immunity due to re-vaccination (some of course, would have pre-existing immunity due to natural infection). In contrast, the UK started a universal influenza vaccination program with the LAIV in 2013-2014 in a few pilot areas. For the 2014-15 season, children aged two to four years and children in school-age were offered LAIV for the first time in some regions of the UK [19], with an estimated coverage ranging from 33 to 41% [49]. Hence, it is expected that children who received the LAIV in the UK had not received an influenza vaccine before, and presumably (because of their young age) had not been exposed to influenza before. We then model two scenarios: 1) a child who has pre-existing immunity due to re-vaccination, identical to the strain contained in the vaccine (so that *d*(*V, P I*) = 0, the American case) and 2) a child who has no pre-existing immunity for whom the vaccine is his/her first exposure to influenza (the British case). Then, we modeled a challenge strain that is 20% different from the vaccine. For the American case, both the vaccine and the challenge strains are contained by pre-existing immunity (Fig. 4A), preventing any B-cell response and a minimal antibody response (Fig. 4C-D), obtaining a negligible within-host vaccine effect (*υe_ω_* = 0.00057). For the British case, the vaccine strain acts as a primary infection (Fig. 4B) eliciting a strong immune response (Fig. 4C and D) that will further control the epidemic challenge, and we obtained a very high within-host vaccine effect (*υe_ω_* = 0.99). If we translate the individual vaccine effects to population-level effects, we would expect to see no vaccine effectiveness in the American case, but to observe vaccine effectiveness in the

**Figure 4:**
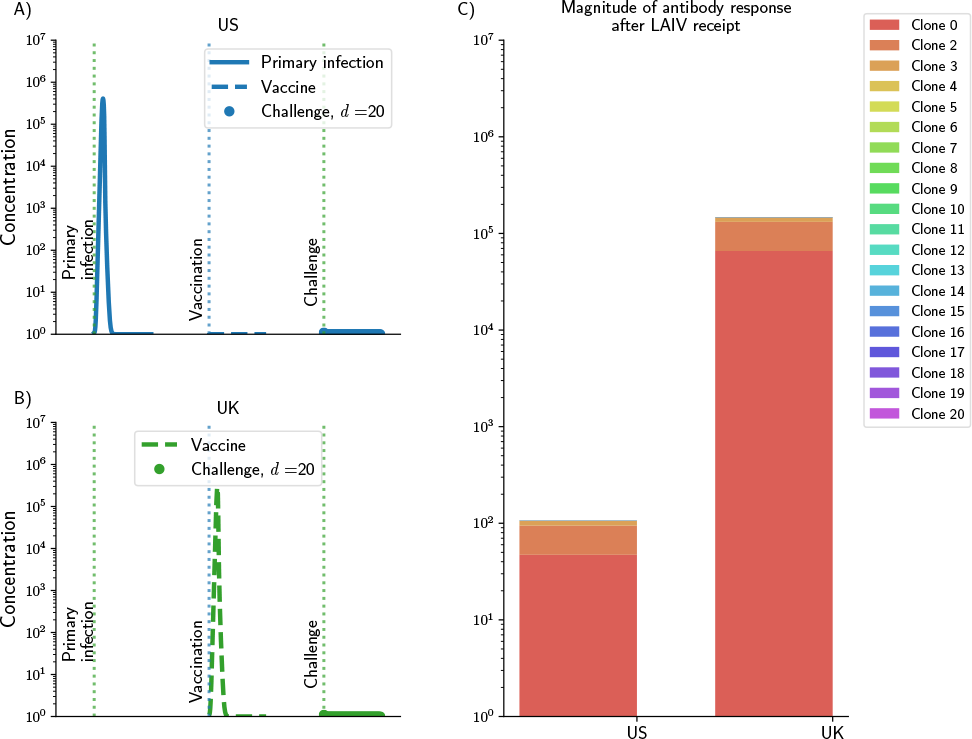
Viral dynamics (panels A and B) and magnitude of the overall antibody response (panel C) for a child in the US (panel A) or in the UK (panel B). For a child in the US, it was assumed that s/he had pre-existing immunity and the strain contained in the vaccine was identical to pre-existing immunity, so that pre-existing immunity controls the vaccine strain and (panels A and C). For a child in the UK, it was assumed that s/he had no pre-existing immunity (panels B and C). In both cases, it was assumed that the challenge strain was 20% different from preexisting immunity, and it is contained by the immune response. However, in the American case, the vaccine is also contained by pre-existing immunity.

## Discussion

Protection due to LAIV has exhibited considerable variability across populations and studies [11, 13, 14, 17–20, 24, 41, 50–53]. In this paper, we use mathematical models to explore a possible hypothesis to explain this phenomenon: interference due to pre-existing immunity. We defined a measure of within-host vaccine effect that allowed us to quantify our results. Furthermore, the results presented here provided us with parsimonious conditions under which the LAIV will be deemed to be efficacious. Our models indicate that there is a window within which LAIV will be successful at controlling a subsequent challenge infection with a drifted virus. This corresponds to the parameter regime where the vaccine is just distant enough from pre-existing immunity that it will replicate and induce an immune response while being close enough to the circulating strain. Otherwise, the vaccine will be inefficacious, either because it is controlled by pre-existing immunity (and this prevents it boosting immunity) or if it is antigenically very distant and stimulates clones too distant from the challenge (a mismatched vaccine). For these cases, the vaccine did not confer any additional protection to the vaccinees, resulting in a poor within-host vaccine effect, and hence we would expect to see a low vaccine effectiveness. There are two additional scenarios that we consider. The first is in individuals such as very young children who have not been exposed to influenza. The second is following the emergence of a pandemic strain of influenza. In both these scenarios there is little effect of pre-existing immunity and a matched vaccine is likely to be efficacious.

Our studies suggest how the current vaccine can best be used, as well as in the choice of strains for inclusion in the vaccine. The current approach using a vaccine strain matched to the circulating strain is likely to be most effective in very young children (who presumably have had little exposure to influenza) and potentially in individuals in areas where there is relatively low attack rate from influenza. Regarding choice of strains for inclusion in the seasonal influenza vaccine – our work suggests that it is important to frequently update the strains as LAIV strains that lag behind the currently circulating strains are likely to have very poor efficacy because of interference with pre-existing immunity. When there is uncertainty in choice of a vaccine strain, our models suggest that the optimal vaccine strain selection is one that is marginally farther away from pre-existing immunity as it is more likely to “take” in a larger fraction of the population. In this regard, our results are consistent with previous results presented by Smith *et al*. for the IIV [29]. Phylogenetic tools such as nextflu [54–58] can provide guidance on which direction to take. In the case of a pandemic we expect less interference with pre-existing immunity and a poorer match with the pandemic strain may well be sufficient.

Our results are also consistent with the current empirical studies and explain the discrepancies observed in the vaccine effectiveness studies done in the US and in the UK for the 2014-15 season. Our results showed that, for a child with pre-existing immunity, as would be a child in the US where children are routinely vaccinated every year, the LAIV would not be immunogenic and would result in a poor within-host vaccine effect. In contrast, we found that in children who are exposed to influenza via vaccination for the first time, as would presumably be the case for a child in the UK, the within-host vaccine effect would be very high. We then hypothesize that the differences in vaccine effectiveness observed on either side of the Atlantic arise from differences in pre-existing immunity. These results point to the need for more studies for LAIV, where it can be fully assessed if the vaccine has triggered an adequate antibody response or it has been inhibited by pre-existing immunity. Such studies are mandatory at the pre-licensure phase of vaccine development, but there are fewer studies done at the post-licensure phase. In an observational study of children aged 5 - 17 years assessing seroconversion and seroprotection for the IIV and the LAIV during the 2013-14 influenza season, King *et al*. found that at most 8% of the participants seroconverted after vaccination with LAIV [8]. In a similar study, Levine *et al* found that only a very small percentage (3-5%) of children seroconverted following receipt of LAIV [59]. Our results suggest that a possible reason for these results can be interference with pre-existing immunity. This is in agreement with previous studies that have shown that pre-existing influenza antibodies can limit both the replication of the strains contained in the LAIV and the immune response generated by the vaccine [60].

Our model, like all models involves a number of simplifications. We focus on the antibody response which is critical to protection against influenza. The antibody-virus affinity was modeled as a percentage of the antigenic distance between the antibody and the virus, and we used the antigenic distance proposed in [29]. By modeling distances in this fashion, we collapsed a multi-dimensional distance into a one-dimensional object, thereby oversimplifying quite complex interactions. Indeed, the antibody-antigen binding process involves not only complementarity in the receptors, but also complementarity in shape, and there are multiple biochemical processes involved (*e.g*. hydrogen bonding). Furthermore, the antigenic distance we used is based on the serum hemagglutination inhibition (HI) assay. This has been historically used to measure the immune response to IIV, but there is increasing evidence that other assays measuring mucosal humoral responses might be better indicators of the LAIV immunogeniticy [61–64]. As more data becomes available on the antibody responses to different epitopes on the hemagglutinin and other proteins of influenza it may be possible to develop strategies that focus on the generation of responses to conserved eptiopes such as those on the stem of hemagglutinin and thus generate a universal influenza vaccine which avoids problems associated with strain variation [65–68]. Another simplification of our model is that we focus on antibody responses and do not consider T cell responses to influenza [69–71]. T cell immunity is largely due to resident memory T cells whose population wanes in the respiratory tract. Integrating both antibody and T cell responses is a direction we hope to work on in the future. Yet another problem is that we have not considered heterogeneity in the responses of different individuals in the population and this may be particularly important when developing multi-scale models that integrate the dynamics of infections of individuals with the spread of the virus in the population.

LAIV is currently being offered to children across the world, however, its effectiveness significantly varies across the years and between countries and studies. LAIV might have advantages over IIV, including a higher acceptance among children, a potential slower waning than inactivated vaccines [72], and higher production capabilities in the event of a new influenza pandemic [73]. Here, we proposed a mechanistic explanation to help understand these differences, and we further propose a way to select the LAIV strain that would have a higher chance of being protective.

## Methods

We developed a series of mathematical models describing the interaction between influenza virus and immune response (SI). We model the adaptive immune response by considering distinct B-cell and antibody clones. The effectiveness of each of these clones depend on the antigenic distance between each clone and the virus. Any antigenic distance based on antibody-virus binding properties is suitable, such as distances based on HI assays (*e.g* [29, 74, 75]) or the *p-epitope* distance [76] provided that it can be transformed into a percentage. For our simulations, we utilized the antigenic distance given in Smith *et al*. [29] and we assumed that once the antibodies and the antigen are at about 20% different (or at distance 4, smiliar to [20]), there is no cross-reactivity. We then transformed this antigenic distance to a match percentage, where *d* = 0%, it is a perfect match, while *d* = 100% implies that the antibody receptors and the virus are totally different (supp Fig A).

We focus here on a target-cell limitation model (**Model 2**, Fig. S1) with 20 B-cell clones. In this model, uninfected cells *U* become infected *I* upon contact with virus *V*. We assume no target-cell production or death given the short timespan of an influenza infection, but infected cells have an increased death rate. In addition, for each clone, B-cells are stimulated and produce antibodies which in turn clear virus. For each clone, B-cell stimulation, antibody production and the antibody’s ability to remove virus are modulated by a function that depends on the antigenic distance between that clone and the virus. Clones closer to the virus will have a rapid expansion while those antigenically different will not be stimulated. We consider two ways in which a vaccine strain can be attenuated: either by reducing its growth rate and/or by increasing the rate at which antibodies remove virus. Table S1 describes the parameters used for this model and presented in the results. A full description of the models and equations can be found in the SI.

We simulated pre-existing immunity by running the model for a “reference” virus, which by simplicity is closest to B-cell clone 1. We then simulated vaccination with a LAIV vaccine by running the model for an attenuated virus (antigen contained in the vaccine). Finally, we simulated an epidemic challenge by running the equations for a third, different virus. We measured the area under the curve (AUC) for the (log) viral load curve for the infection with the challenge strain and use this as an indication of immune control: if the *AUC <* 2 the infection with the challenge virus was deemed controlled. Furthermore, we use this quantity to define a within-host vaccine effect *υe_ω_*, that measures the ability of the vaccine to prevent further infections with that particular challenge strain, given by 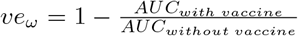. A full description of the methods can be found in the SI. We performed a sensitivity analysis (SI) and showed that our results were consistent for all the models proposed and for a wide range of parameter values.

## Supporting Information (SI)

### Mathematical models

In this section we present four models of increasing complexity considered in the present work. For each model, we divide the B-cell population into *n* discrete clones, with different clones having different affinities to different viruses. Conceptually, the effectiveness of antibody-binding and the stimulation of B-cells depend on the antigenic distance between the virus and the B-cells, and we model it through a Hill function given by

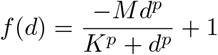

where *M*, *K*, and *p* are parameters to be determined. The variable *d* measures the distance between antibodies *A* and virus *V* as a percentage of a preselected antigenic distance. Any antigenic distance based on antibody-virus binding properties is suitable, such as distances based on HI assays (*e.g* [29, 74, 75]) or the *p-epitope* distance [76] provided that it can be transformed into a percentage, where *d* = 0%, it is a perfect match, while *d* = 100% implies that the antibody and the virus are totally different. The models are based on earlier models [42, 77–79], and described in more detail below.

**Model 1:** The simplest possible model of interactions between influenza viruses and the immune system consist of a model involving virus *V* and B-cells. B-cells proliferate at a rate *σ* depending on the concentration of virus. The B-cell population is divided into *n* distinct clones, where each clone *B_i_* (*i* = 1,…, *n*) has a different efficacy to eliminate virus depending on their antigenic distance *d_i_* to it. For each clone, the function *f*(*d_i_*) described above modulates the ability of that clone to control the virus, as a function of the distance *d_i_* in two different ways: 1) affecting the amount of B-cells produced and 2) affecting the killing rate of virus by B-cells. Virus grows at a rate *r* and is eliminated by B-cells at a rate *k*. With these hypothesis, we obtain the following system of ordinary differential equations

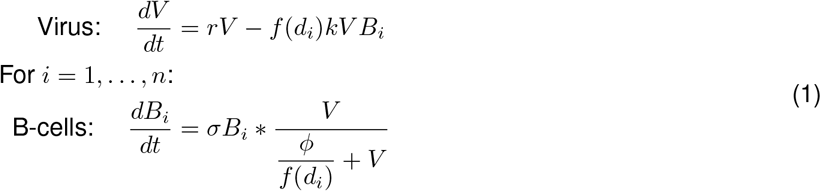

**Model 2:** We next add uninfected cells *U*, infected cells *I*, and antibodies to our model, obtaining a target-cell limitation model, similar to the classic viral kinetic model ([80, 81]. Uninfected cells get infected upon encounter with virus with infectivity *β*. The full description of this model can be found in the main text. The corresponding ordinary differential equations are given by

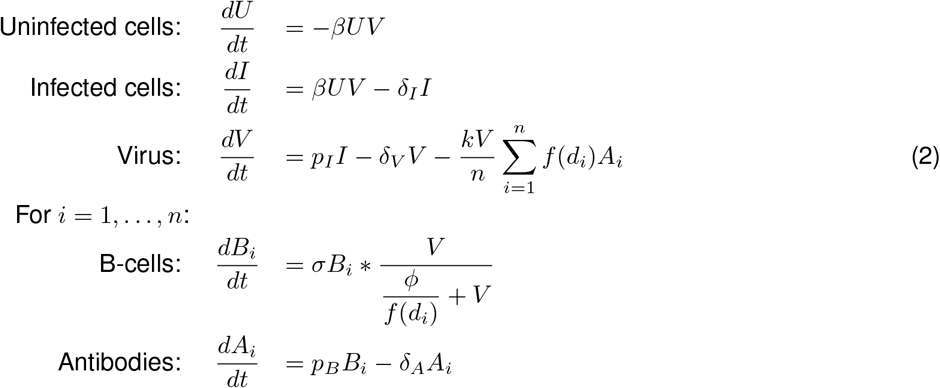

**Model 3:** This model builds upon the last one by adding innate immunity. Innate immunity in this model causes uninfected cells to become refractory to infection. This is modeled by the removal of cells from the *U* population at a rate proportional to the amount of innate immunity (which is proportional to the number of infected cells).

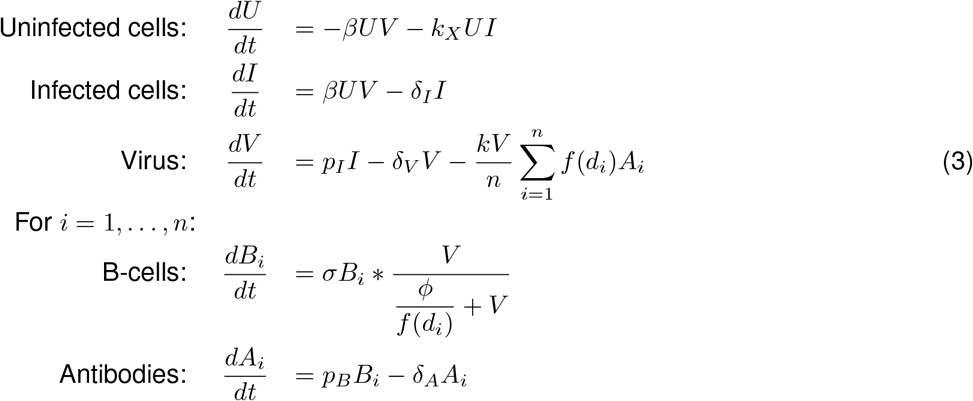

**Model 4:** In this model we add an equation to explicitly model innate immunity *X*. Innate immunity grows in a logistic fashion at rate *σ_X_*, and it is modulated by the amount of virus present in the system via a saturation function, where *ϕ_X_* represents the half-saturation constant. Finally, innate immunity decays at a rate *δ_X_*,

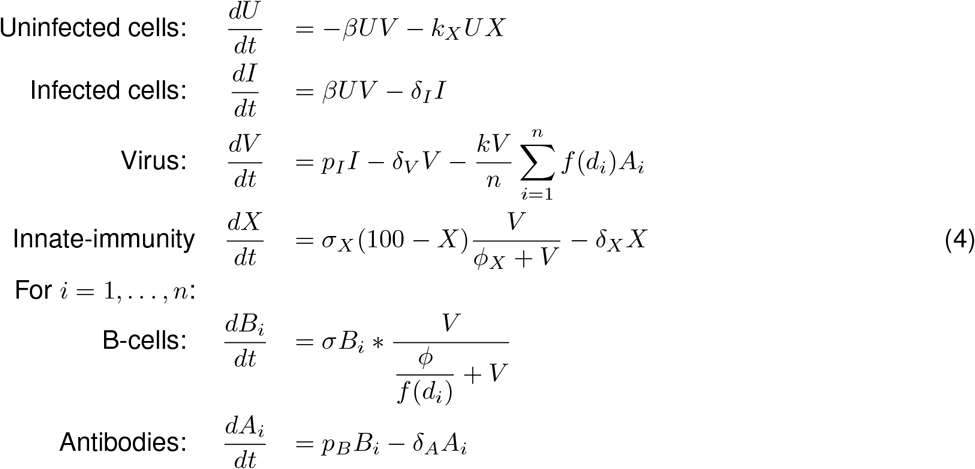

### Sensitivity analysis

As in the main text, we denote the distances between vaccine and pre-existing immunity, vaccine and challenge, and challenge and pre-existing immunity by *d*(*V, P I*), *d*(*V, C*) and *d*(*C, P I*) respectively. We performed sensitivity analysis by varying our models and our parameters. Our results were consistent over a variety of parameter values for all the models considered. The different assumptions leading to our different models did not qualitatively alter our results, but, as expected, they altered the shape and particular range of values for which the conclusions hold. We varied the cross reactivity of antibodies by varying the antibody affinity to virus so that they stop being reactive to virus once their antigenic distance to virus was 15% to 40% (figs. S4, S5, S6, S7). When the antibodies are less cross-reactive, then as expected, vaccine and pre-existingn immunity stop being protective when their distance to the challenge is smaller (*d*(*V, P I*) ~ 10%). Under this scenario, there is a wider range of values for which choosing a vaccine strain farther away from pre-existing immunity would result in a better control of subsequent challenges (Fig. S4). The area for which the within-host vaccine effect is high follows the diagonal (so the vaccine will be highly efficacious for vaccines matched to the epidemic strain if both are sufficiently different from pre-existing immunity), see Fig. S6. When the antibodies are more cross reactive, our models predict that pre-existing immunity will control the epidemic challenge and the vaccine strain for a wider range of antigenic distances, resulting in a smaller range where the vaccine will be immunogenic (Fig. S5 and Fig. S7).

**Figure S1:**
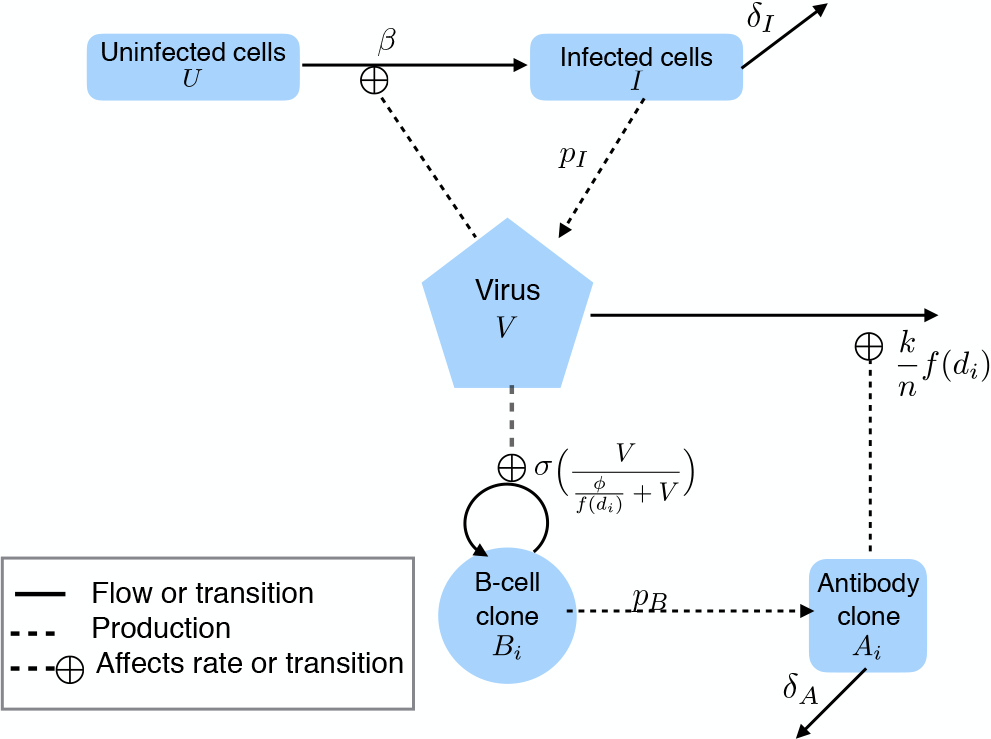
Diagram of Model 2 (equations and full description can be found in the SI). Uninfected cells *U* become infected *I* upon contact with virus *V*. Infected cells produce virus at rate *p_I_* and are removed at rate*δ _I_*. B-cells are produced at a rate *σ*, They are further divided into *n* distinct clones and produce antibodies at rate *p_B_*. Antibodies are removed at rate *δ _A_* and eliminate virus at a rate proportional to their antigenic distance to the virus (*d_i_* with *i* = 1,…, *n*), determined by the function *f*(*d_i_*).

**Figure S2:**
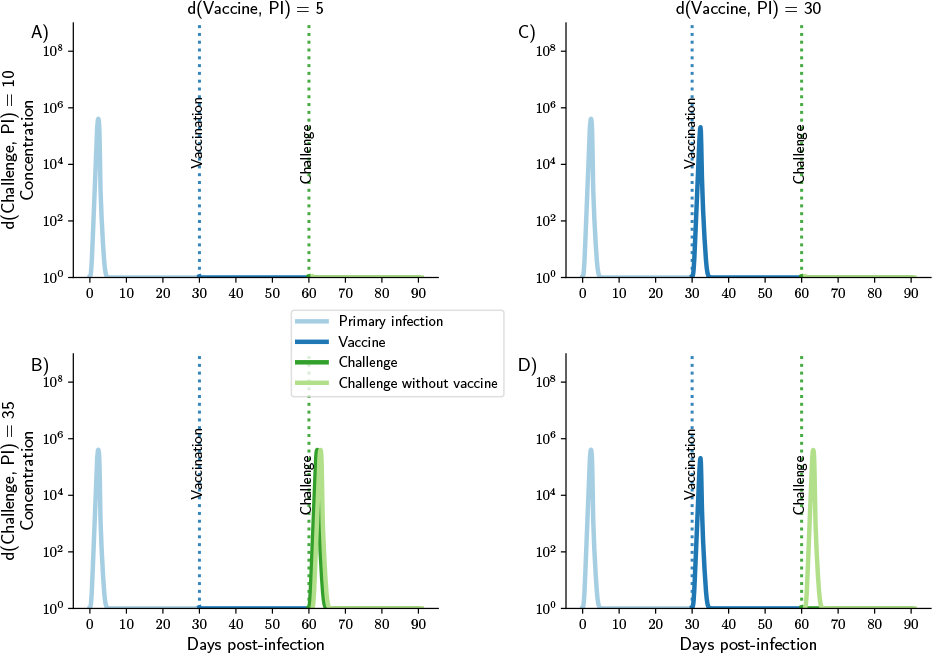
Viral load curves for 4 different scenarios depending on the antigenic distance between pre-existing immunity, vaccine strain and challenge strain. Top row: challenge strain is 10% different from pre-existing immunity, bottom row: challenge strain is 35% different from pre-existing immunity with vaccine 5% different from pre-existing immunity (panels A and C) or 30% different (panels B and D). Distance between vaccine and challenge: 5%, 20%, 30% and 5% respectively for panels A-D

**Figure S3:**
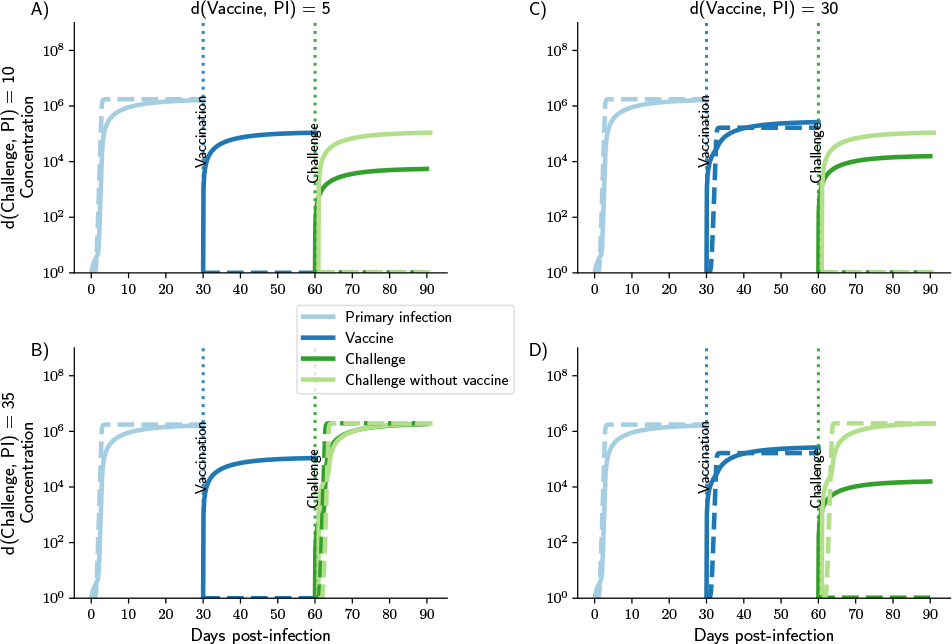
Total B-cell (dashed lines) and antibody (solid lines) dynamics for 4 different scenarios depending on the antigenic distance between pre-existing immunity, vaccine strain and challenge strain. Top row: challenge strain is 10% different from pre-existing immunity, bottom row: challenge strain is 35% different from pre-existing immunity with vaccine 5% different from pre-existing immunity (panels A and C) or 30% different (panels B and D). Distance between vaccine and challenge: 5%, 20%, 30% and 5% respectively for panels A-D

**Figure S4:**
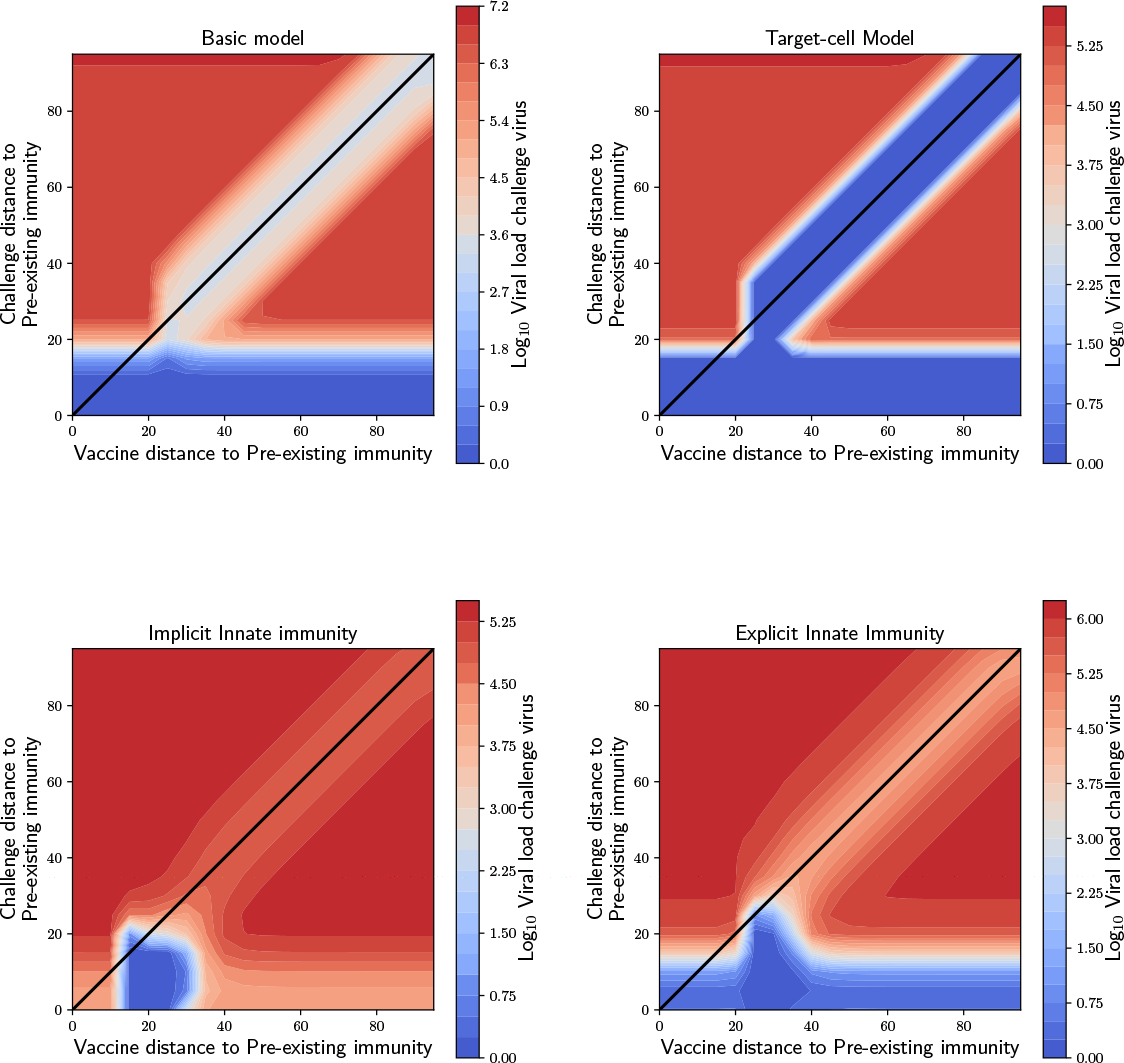
Sensitivity analysis for the contour plots representing the severity of infection presented in the main text when the antibodies are less cross-reactive. Here, the parameters for the Hill function were *K* = 8, *p* = 10. This results in antibodies that stop being reactive to a virus once their antigenic distance differs by ~15%.

**Figure S5:**
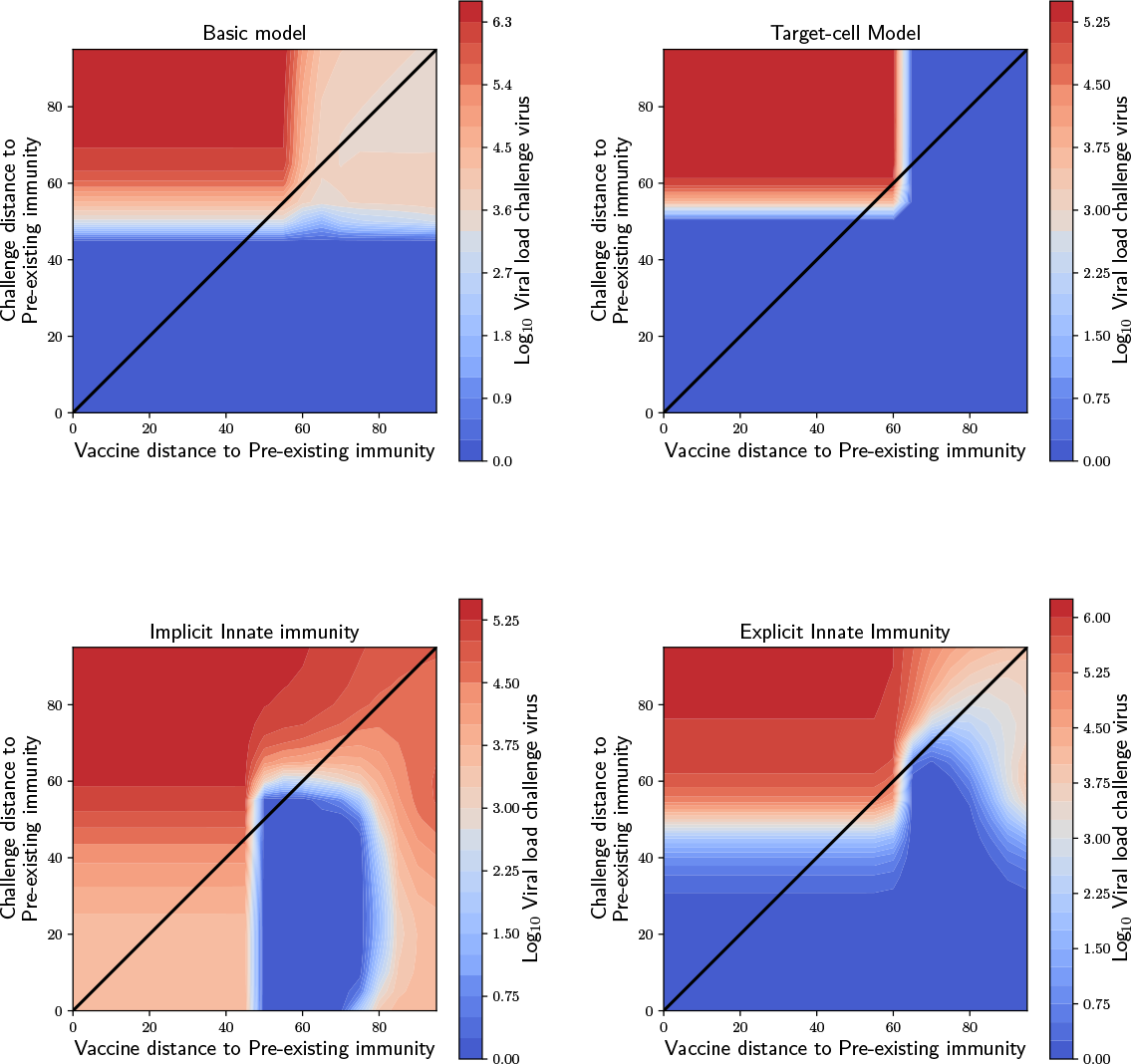
Sensitivity analysis for the contour plots representing the severity of infection presented in the main text when the antibodies are more cross-reactive. Here, the parameters for the Hill function were *K* = 25, *p* = 15. This results in antibodies that stop being reactive to a virus once their antigenic distance differs by ~40%.

**Figure S6:**
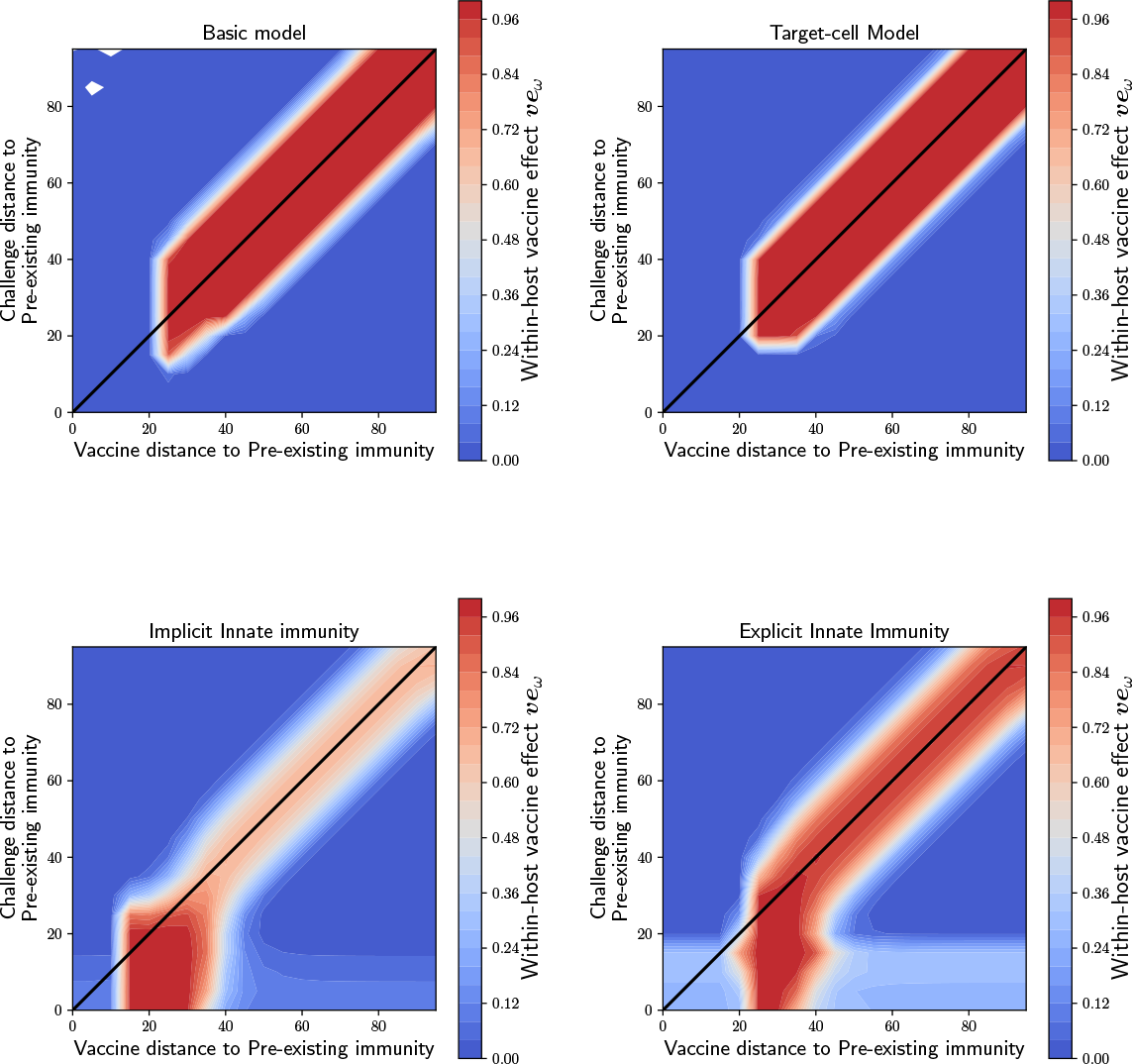
Sensitivity analysis for the within-host vaccine effect contour plots presented in the main text when the antibodies are less cross-reactive. Here, the parameters for the Hill function were *K* = 8, *p* = 10. This results in antibodies that stop being reactive to a virus once their antigenic distance differs by ~15%.

**Figure S7:**
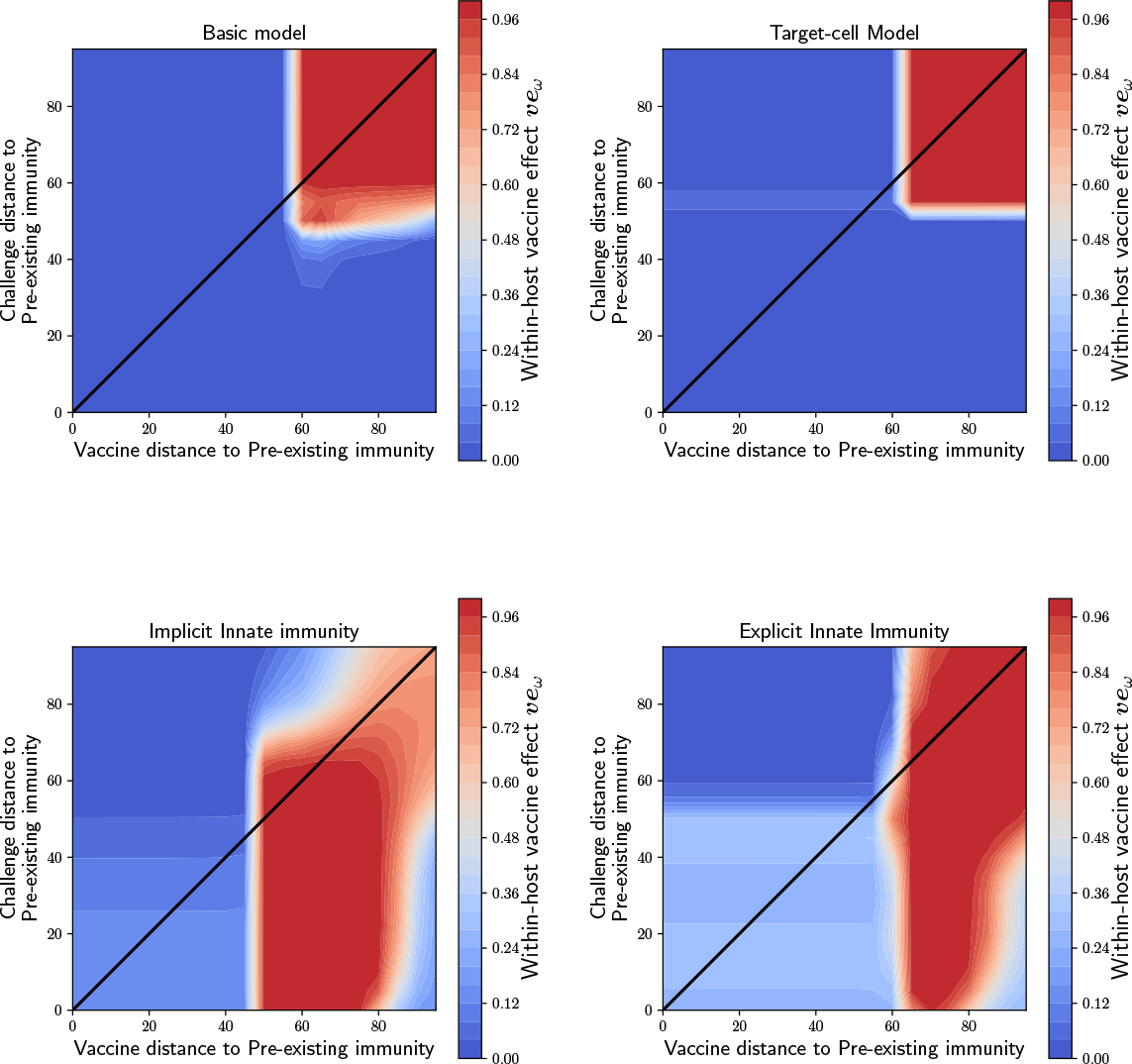
Sensitivity analysis for the within-host vaccine effect contour plots presented in the main text when the antibodies are more cross-reactive. Here, the parameters for the Hill function were *K* = 25, *p* = 15. This results in antibodies that stop being reactive to a virus once their antigenic distance differs by ~40%.

**Table S1:** Model 2 used in the main text: parameters definitions and values.

| Model parameter | Symbol | Units | Value | Reference |
| --- | --- | --- | --- | --- |
| Virus infection rate | $\beta$ | | $2.7 * 10^{-5}$ | [42] |
| Virus infection rate for vaccine strain | $\beta_{Vac}$ | | $(0.1 - 1) * \beta$ | varied |
| Death rate for infected cells | $\delta_I$ | day<sup>-1</sup> | 4.0 | [42] |
| Virus production rate | $p_I$ | day<sup>-1</sup> | $1.2 * 10^{-2}$ | [42] |
| Antibody death rate | $\delta_A$ | day<sup>-1</sup> | $10^{-1}$ | [82] |
| Virus clearance rate | $\delta_V$ | day<sup>-1</sup> | 3 | [42] |
| Infected cell killing rate by antibodies | $k$ | day<sup>-1</sup> | $5 * 10^{-3}$ | |
| Infected cell killing rate by antibodies for vaccine virus | $k_{Vac}$ | day<sup>-1</sup> | $(10 - 50) * k$ | varied |
| Number of virus for half-max. B-cell activation | $\phi$ | | $5 * 10^3$ | scaled <sup>a</sup> |
| Max. activation rate of B-cells | $\sigma_B$ | | 10 | scaled |
| Rate of production of antibodies by B-cells | $p_B$ | day<sup>-1</sup> | $10^{-1}$ | scaled |

<sup>a</sup>These parameters were scaled so that the B-cells and antibodies were at equilibrium when no infection is present.

**Table S2:**
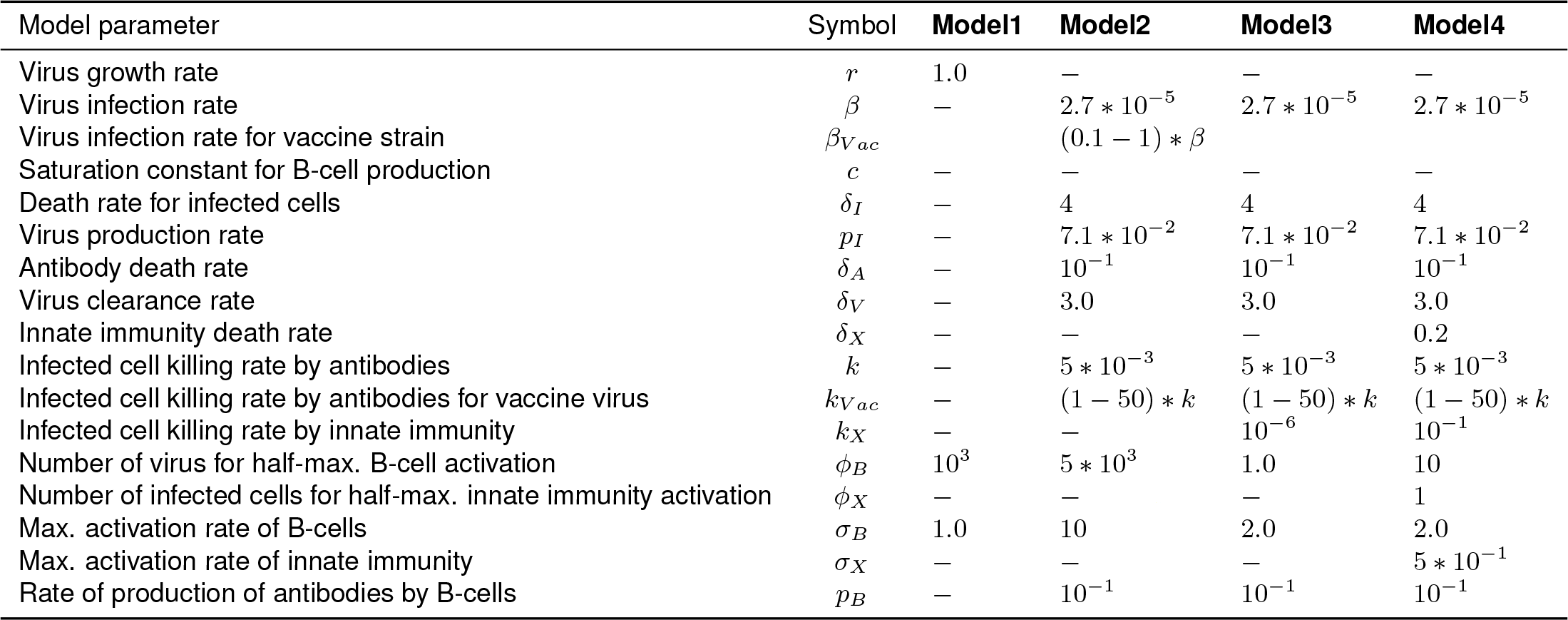
Parameters for all models.

